# Landing on a swinging perch: peach-faced lovebirds prefer extremes

**DOI:** 10.1101/2024.07.21.604435

**Authors:** Partha S Bhagavatula, Andrew A. Biewener

**Affiliations:** Concord Field Station, Department of Organismic and Evolutionary Biology, Harvard University

**Author notes:** Correspondence: Partha S Bhagavatula Ph.D. OEB Associate, Concord Field Station, Harvard University, 100 Old Causeway Rd. Bedford, MA 01742 USA.

**Keywords:** Peach-faced lovebird, aerial landing, moving perch, kinematics, force, torque

## Abstract

Birds frequently must land safely and accurately on moving branches or power lines, and seemingly accomplish such maneuvers with acrobatic precision. To examine how birds target and land successfully on moving supports, we investigated how peach-faced lovebirds (*Agapornis roseicollis*) approach and land on a swinging perch. Lovebirds were trained to take off from a hand-held perch and fly ∼6 m to land on a servo-controlled swinging perch, driven at three sinusoidal frequencies, in a purpose-built flight corridor. Lovebird flight and landing kinematics were recorded using a motion capture system. A force-torque sensor mounted to the landing perch recorded the bird’s horizontal and vertical landing force and pitch torque. In support of our hypothesis for stable landings, lovebirds timed their landings in a majority of trials (51.3%), when the perch was approaching either extreme of its motion with its velocity nearing zero (27.5% in the same direction as the bird’s approach – SDs, and 23.8% in the opposite direction to the bird’s approach – ODs). As a result, lovebirds exhibited a robust bimodal strategy for timing their landing to the phase of the swinging perch. Less commonly, lovebirds landed when the perch was moving at high velocity either toward the bird’s approach (12.3%) or in the same direction as the bird’s approach (11.5%); with the remainder (21.9%) of trials distributed over a broad range of swing phases. Landing forces were greatest in the horizontal plane, with vertical forces more varied and of smaller magnitude across all landing conditions. This reflected the shallow flight trajectory (center of mass approach angle: −13.2 ± 3.0o SEM relative to horizontal) that the lovebirds adopted to decelerate and land. Increased landing force correlated with greater landing speed of the bird relative to the perch (R2 = 0.4296, *p <* 0.0001). The lovebirds initiated landing with a consistent body pitch angle (81.9 ± 0.46o SEM relative to horizontal) across all landing conditions, using the horizontal perch reaction force to assist in braking when landing. Correspondingly, the landing angle of the feet relative to the perch support was 56.9 ± 2.8o. Subsequent head-down body pitch rotation of the bird after landing was not well correlated and generally opposite to the initial direction and magnitude of landing pitch torque, which was generally negative due to foot rotation and ankle flexion at landing. Flexion of the birds’ hind limb joints (ankle: - 29.2 ± 9.2o, knee: −13.6 ± 7.4o, and hip: −4.0 ± 3.4o at landing, combined with their horizontal approach trajectory, reduced the magnitude of landing torque by aligning the bird’s center of mass trajectory more closely to the landing perch (3.61 ± 0.21 cm) than if they landed from above the perch. Landing pitch torque and body pitch rotation also increased uniformly in response to increased perch swing frequency. In contrast to landing forces, landing pitch torque was more varied across landing conditions, as well as in relation to the phase of landing. In general, higher landing force was encountered when the perch was moving towards the approaching bird. Our results indicate that lovebirds regulate their approach trajectory and velocity to time the phase of landing to a moving perch, providing insight for designing biologically-inspired unmanned aerial vehicles capable of landing on moving targets.

## Introduction

Volant animals select and target landing sites that are often unstable and moving in complex ways. Much work has been focused on how birds (e.g. pigeons and hawks) use visual cues to land on fixed, stationary targets (1–4). However, despite their ecological relevance, the strategies that flying animals use to land on moving targets has been less well studied (5).

Further, relatively few studies have focused on the kinematics and biomechanics of landing by flying and gliding vertebrates. Studies of take-off and landing flights of starlings (*Sturnus vulgaris*) (6), pigeons (*Columba livia*) (7), diamond doves, and zebra finches (8, 9) between fixed perches have shown that take-off forces exceed landing forces and that pigeons, as well as diamond doves and zebra finches, change the their tail angle and wing stroke-plane angle by altering overall body pitch, pitching up at landing to increase drag and slow the bird’s landing velocity. In their study of diamond doves and zebra finches, Provini et al. (2014) used particle image velocimetry to show that the change in body pitch of diamond doves and zebra finches when landing alters the induced downwash produced by the wings from a vertical to nearly horizontal orientation, consistent with a change in the wings’ angle of attack, increasing drag to assist in deceleration of the bird prior to landing.

Studies of gliding vertebrates – northern gliding squirrels, *Glaucomys sabrinu*s (10) and gliding lizards, *Draco dussumieri* (11) – also show that gliders pitch-up to decelerate when landing to a stationary perch. Finally, a study of three species of bats landing to a ceiling-mounted force platform (12) also undergo pitch rotations (as well as body roll) to land, and revealed that a foliage-roosting species (*Cynopterus brachyotis*) used four-point landings that incurred much higher body weight-normalized landing forces than two cave-dwelling species (*Carollia perspicillata* and *Glossophaga soricine*) that used their hindlimbs for two-point landings.

Here, we examine how peach-faced lovebirds (*Agapornis roseicollis*) target and stabilize landings on a swinging perch, emulating a tree branch or a telegraph wire moved by the wind. We ask, how are landings timed by lovebirds to improve landing stability and safety in relation to the magnitude of landing forces, pitch torque and body rotations that they experience?

When landing, flying animals must use visual cues to accurately locate a suitable perch relative to their own motion and then land successfully by braking to stabilize their body’s motion when their feet contact the perch. Past work has shown that pigeons may use motion parallax cues to target a perch by bobbing their heads during a landing approach (13). Head bobbing may serve to help disambiguate the target’s location relative to the optic flow of surrounding features generated by the animal’s own self-motion. However, in a recent study of lovebirds (5) landing on a stationary perch allowed to swing in reaction to the bird’s landing, no evidence of head bobbing was observed.

In addition to visually locating a suitable landing target, flying animals must also achieve stable landings by controlling the forces and torques transmitted to their legs and feet after contacting the landing surface. Recent work (14) has shown that Pacific parrotlets achieve robust gripping forces by adjusting the dynamics of their feet, toes and claws when landing on stationary perches of varying texture and diameter. Achieving a mechanically stable landing is likely even more of a challenge when a perch is moving. By recording the kinematics and landing kinetics of lovebirds trained to fly to a sinusoidally swinging perch, we seek to evaluate how lovebirds adjust their approach speed and time their landings relative to the perch’s motion. Native to southern Africa, peach-faced lovebirds possess a well-developed visual system (15), with laterally positioned eyes that, similar to most bird species, have a relatively small extent of binocular overlap (16). They live in a seasonally changing environment composed of dry open country, wooded grassland and trees (15); consequently, lovebirds experience a diverse set of landing sites within their natural habitat. Given this, they serve as an attractive model system to investigate the biomechanics and landing behavior of birds to moving perches.

To maximize their control for stable landings, we hypothesize that lovebirds will exhibit a preference for landing phases relative to the perch’s motion that minimizes their body pitch about the perch and contact reaction forces. Accordingly, we test the following specific hypotheses for the landing dynamics associated with different phases of landing on a sinusoidally swinging perch (Fig. 2).

We hypothesize that when landing at or near the extremes of the perch’s motion (either same direction slow – **SDs**, or opposite direction slow – **ODs**), minimum pitch torque (+/-**TY**,min) and landing forces will be experienced, similar to the torque and forces experience when landing on a stationary perch (zero-velocity, **Vo**, control landing: **So**)

However, when landing as the perch approaches zero velocity (**V∼o**) in the same direction as the bird’s approach (**SDs**) at the end of its swing (Fig. 2C; or when re-accelerating to begin its swing), we hypothesize landing will result in a negative (head-up) pitch torque to achieve a stable landing. By contrast, when the perch is moving slowly in the opposite direction to the bird’s approach (**ODs**, Fig. 2D), the bird will generate a positive (head-down) pitch torque to stabilize its landing. Landings to a stationary perch (**So**) will result in variable patterns of negative or positive pitch torque (+/-**TY**,min).

Correspondingly, we hypothesize that lovebirds will minimize the frequency of landings when the perch is moving at its fastest speed (mid-swing) either in the same direction (**SDf**, Fig. 2E) as the bird’s approach or in the opposite direction (**ODf**, Fig. 2F). Landing when the perch is moving in the same direction to the bird’s approach (**SDf**) at peak velocity (**+Vmax**) mid-swing will result in a strong rearward (head-up) pitch torque (-**TY**,min); whereas, landing when the perch is moving in the opposite direction to the bird’s approach mid-swing will result in a strong forward (head-down) pitch torque (+**TY**,min) of the bird. In either case, pitch moments will initially be high before the bird stabilizes its body on the swinging perch. As a result, we hypothesize that angular pitch rotations of the bird upon landing will correlate with the magnitude and direction of pitch torque exerted on the perch.

Finally, we hypothesize the landing forces will vary with the relative speed of the bird to the motion of the perch, such that a larger relative landing speed will generate greater perch landing forces acting on the bird’s legs and body to decelerate its landing momentum.

## Methods

### Ethics statement, subjects, and housing

All experiments were carried out with the approval of the Faculty of Arts and Sciences IACUC of Harvard University, Cambridge, MA (AEP 98-04). Four wildtype Peach-faced lovebirds (body mass: mean 51.7 ± SD 4.9 g = 0.53 N mean body weight, BW) were housed communally in an aviary (12:12hr L:D cycle) with perches and toys for enrichment at the Concord Field Station, Harvard University (Bedford, MA, USA). Birds were fed commercial birdseed (Roudy Bush Maintenance, Mini, Scott Pharma, Inc, USA) comprising essential minerals and enriched with vitamins.

### Flight corridor, perch control and training

A custom-made flight corridor (Dimensions: Length 6.02 m, Width 1.36 m and Height 1.22 m) constructed of plywood was installed indoors for the experiment, with a 1.27 cm mesh nylon netting cover (Fig. 1). The corridor walls were painted white and decorated with vertical stripes (a square wave grating) made of black paper (Jet Black A1, Code: 402275004, Canford paper, Daler Rowney, England) 10 cm wide, spaced 10 cm apart. The landing perch consisted of black painted wooden dowel, of length 38.5 cm and diameter 8.77 mm suspended by a tubular aluminum and carbon fiber frame held in place by 3D printed plastic fixtures, supported between a digital servomotor (HS-805 MG, Hitec RCD, USA) and a bearing mount. The digital servomotor was controlled by a USB Servo Controller (Micro Maestro 6-Channel (Assembled), Pololu, USA). The entire perch assembly was suspended from a ‘T’-slotted aluminum frame (80/20 Inc, USA). A 6 DOF force-torque sensor (Nano 43, ATI Industrial Automation, USA) supported the landing perch. The corridor was devoid of visual landmarks other than the perch, the side and end walls. The black perch offered high contrast against the white background wall. The floor was covered with white butcher paper to improve contrast for filming the bird from above.

**Figure 1.**
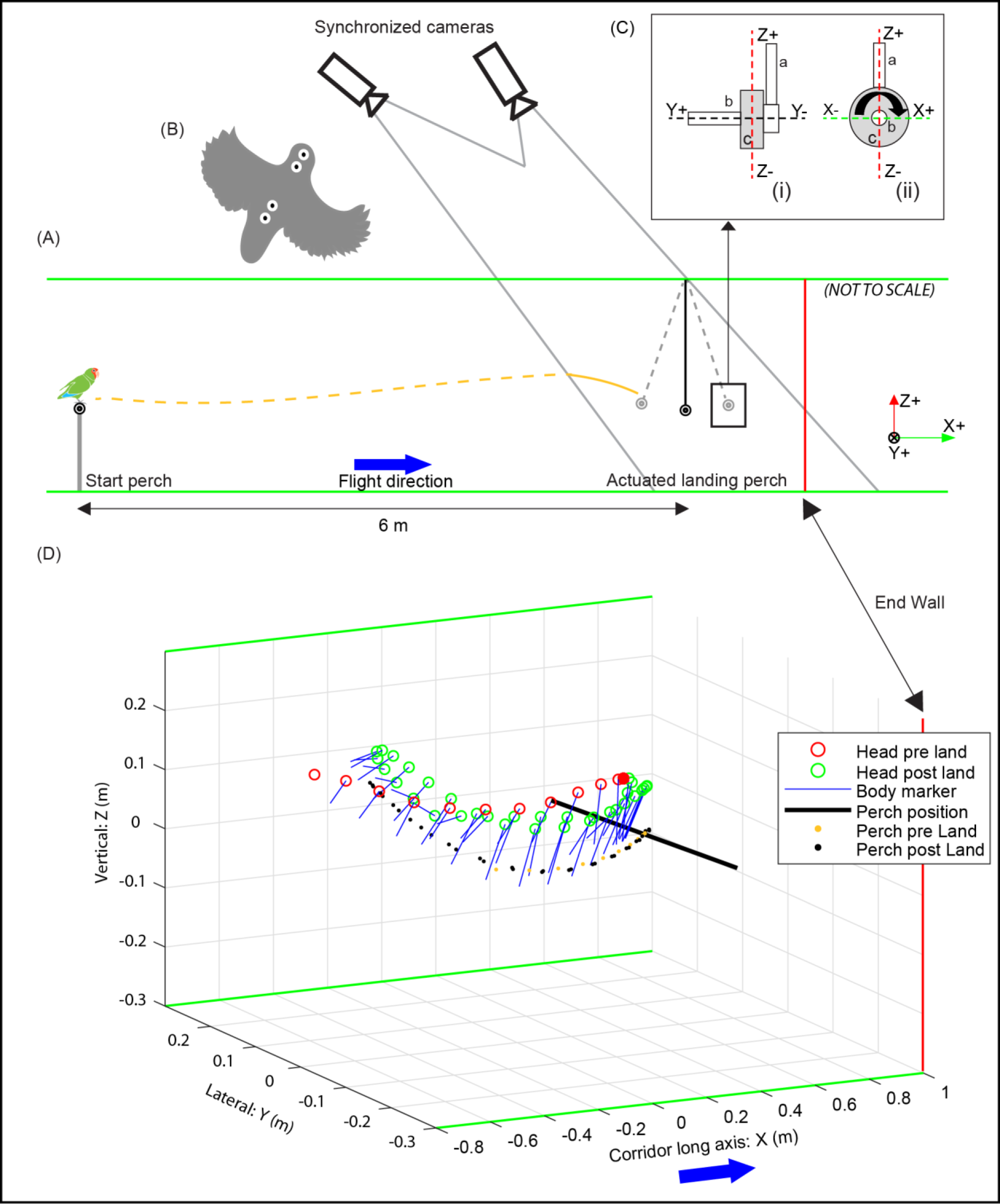
Experimental setup for lovebird landings on a moving perch. (A) Starting from a takeoff perch, birds flew six meters and landed on a swinging (or stationary) perch. (B) Tracked marker positions on the dorsal side of the bird’s head and body. (C) Load cell (c) shown in light grey, attached to the end of perch (b) and supported by a lightweight aluminum rod (a) connected to a servomotor at the top end (not shown), in front (i) and side (ii) views, with XYZ axes defined as for flight corridor. Also not shown was a Mobius (Model: 1S) camera mounted on the opposite end of the perch frame support that provided close-up kinematic recordings of the birds’ landings. The curved solid black arrow denotes + head down pitch torque (**T_Y_**) about the perch axis. (D) Shows a representative landing on the actuated perch. Red circle: head position pre-landing; green circle: head position post-landing; solid red circle: head position at the time of landing; blue line shows body orientation; horizontal black line: indicates perch position at landing; yellow dotted line: perch’s position pre landing; black dotted line: perch’s position post landing. Green lines (A and D): corridor boundaries. Red lines (A and D): corridor end wall. A solid blue arrow shows the direction of bird flight (A and D). Flight corridor and load cell axes (A and D): X – corridor long axis, Y – lateral, Z – vertical.

The digital servo was controlled by means of a servo control software: Maestro Control Center (Pololu, USA). The software allows for precise and smooth control of the servo movement by controlling its speed and acceleration, which enabled control of the perch at three angular frequencies (0.21, 0.49, 0.62 Hz), in addition to being held in a stationary position (control) with the servo-motor on (see Table 1). However, in order to minimize vibrations of the perch induced by its sinusoidal motion, perch amplitude was increased at the highest perch frequency (23.9o @ 0.62 Hz versus 12.2o @ 0.21 Hz and 16.5o @ 0.49 Hz).

**Table 1:** Results of Hartigan’s Dip statistic and Gaussian Goodness of fit to determine the type of bird landing distribution. (n= 400). SSE: Sum of Squared Errors.

| Test name | Same Direction | Test result | Opposite Direction | Test result |
| --- | --- | --- | --- | --- |
| **Hartigan's Dip statistic** | Dip = 0.1159<br>$p < 0.0001$ | Multimodal distribution | Dip = 0.1290<br>$p < 0.0001$ | Multimodal distribution |
| **Gaussian Goodness of fit** | SSE: 149.1<br>$R^2 = 0.9199$<br>$p < 0.0001$ | Multimodal distribution | SSE: 157.7<br>$R^2 = 0.8933$<br>$p < 0.0001$ | Bimodal distribution |

The lovebirds were trained to hold their position on a hand-held perch and to take off upon slow rotation of the perch to fly down the corridor and land initially on a stationary perch, and subsequently on a sinusoidal swinging perch. For every bird, this molding and training practice required around 30 flights, spaced over three days. Once training was complete, birds were flown under different experimental conditions, which involved landing on an actuated perch while being video-recorded to obtain kinematics data. For each bird, 25 trials were recorded for each of the three sinusoidal frequencies and for comparison to landings to a static perch that served as a control. Flights were randomised across these four conditions (n = 100 trials per bird, total n = 400 trials, N = 4 birds). Each bird was marked using white circle of latex (Liquid paperTM, USA) with a black permanent marker center dot on feathers at four locations: two on the head (forward and rear head marker) and two on the body (dorsal thorax and rump marker) for kinematic tracking as shown in Figure 1B. The wing tips were not marked and were manually tracked against a high contrast white background.

### Filming of flights

Flights were captured using two black and white high-speed (200 Hz), high-resolution video cameras (Model N5S1, Integrated Design Tools Inc, USA; 2336 X 1728 pixel images) attached to lenses (AF Zoom Nikkor 24-85mm f/2.8-4D IF, Japan) placed obliquely to cover the full envelope of perch displacement and a small region (∼1.8 m) from the direction of bird approach towards the perch. The high-speed cameras were controlled using Motion Studio video acquisition software (Integrated Design Tools Inc, USA). Four 750 W halogen lamps (Lowel Tota, Tiffen, NY, USA) were used to light the landing area.

In addition, a MobiusTM action camera (Model: S1, Huizhou tuopu xunshi Technology Co., Ltd, PR China) operating at 60 fps was mounted on the left lateral perch support to provide a zoomed in lateral view of the lovebirds, as they landed on the perch. The videos of three of four birds (the fourth landed too close to the camera, yielding out-of-focus and blurred recordings) were analyzed to evaluate the landing kinematics of the feet and hind limbs relative to the bird’s trunk, as well as the bird’s feet and center of mass (COM) approach trajectories relative to perch, as it landed. To estimate the bird’s COM, a frozen lovebird with wings depressed against the body was suspended by 3-0 silk suture ties from three locations on its trunk yielding the center of gravity intersection as the COM. With the wings manipulated into an elevated position characteristic of landing (see suppl. Movies 1-4), the bird refrozen and suspended, the COM shifted by < 1.2 mm, which was deemed insignificant with respect to estimating COM landing trajectory. These recordings also enabled us to quantify the body pitch angle offset (Θ2, Fig. 2), measured from the bird’s head to the midpoint of the tail base relative to that measured by the IDT cameras from above, based on the dorsal head and body markers (Θ1). Body pitch angle was then quantified as the sum of Θ1 + Θ2.

**Figure 2.**
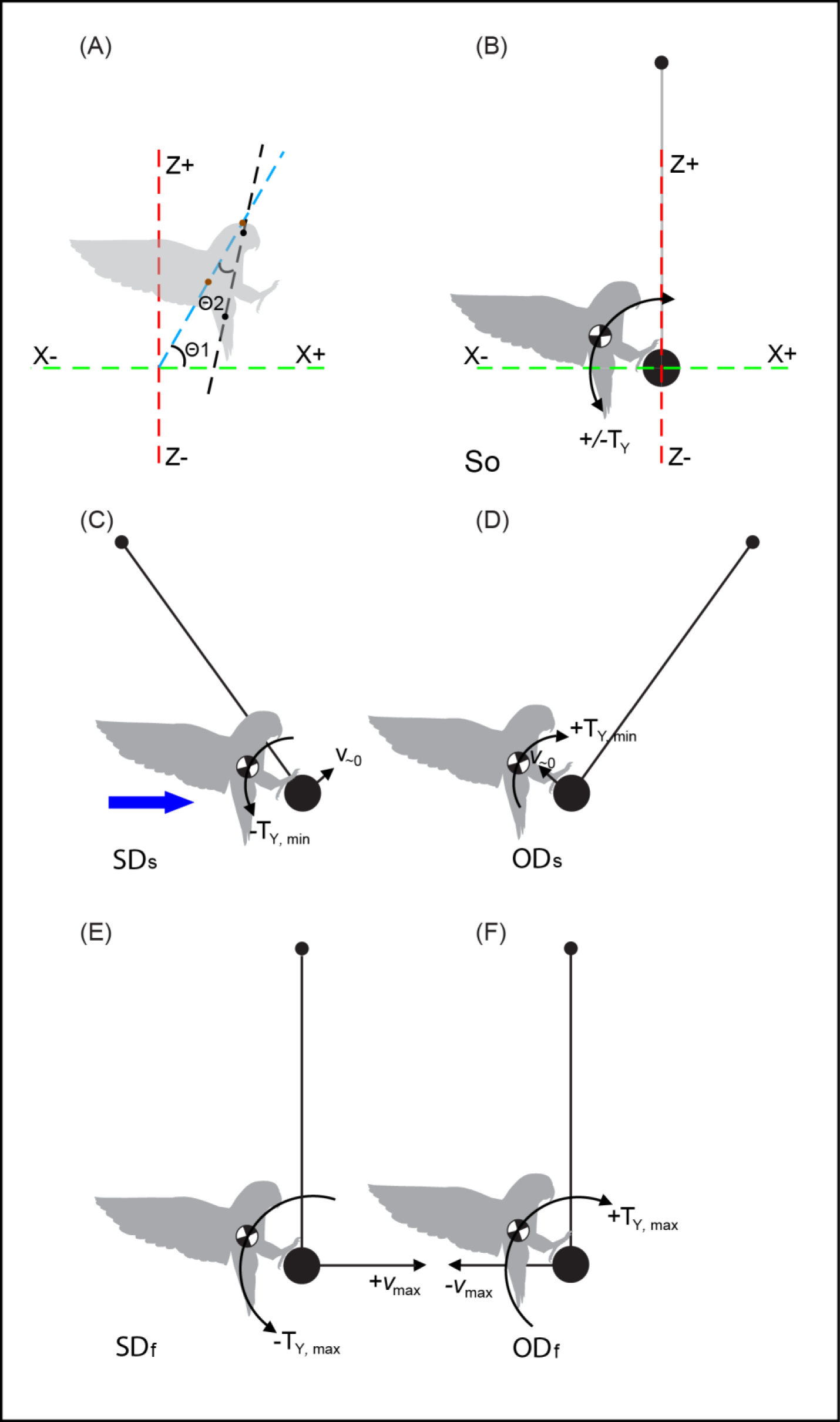
Pitch angle and hypothesized lovebird landing dynamics for different landing conditions. **(A)** Body pitch angle (**Θ = Θ1+ Θ2**) subtended by a tangent connecting the head and dorsal body marker (blue dashed line, **Θ1**) determined from the IDT cameras and body pitch offset (black dashed line, **Θ2**) determined from the Mobius camera recordings (see methods), relative to global horizontal (green dashed line). A solid blue arrow shows direction of bird flight. Hypotheses for landing conditions: **(B)** Hypothesis 1: minimum pitch torque (+/-**TY**,min) exerted on a stationary perch (**So**); **(C)** Hypothesis 2: landing in same direction with respect to the perch’s motion when the perch approaches zero velocity (**V∼o**) (**SDs** – same direction slow), minimum head-up (negative) pitch torque (-**TY**,min); **(D**) Hypothesis 3: Landing in opposite direction relative to the perch’s motion as perch approaches zero-velocity (**ODs** - opposite direction slow) minimum initial head-down (positive) pitch torque (+**TY**,min); **(E)** Hypothesis 4: landing in same direction as the perch’s motion when perch approaches maximum velocity (**+Vmax**) (**SDf** - same direction fast), maximum negative pitch torque (-**TY**,max); **(F)** Hypothesis 5: landing in opposite direction to the perch’s motion at maximum perch velocity (**-Vmax**) (**ODf** - opposite direction fast) maximum positive pitch torque (+**TY**,max). A range of perch-velocity conditions exist between **(C)**, **(D)**, **(E)** & **(F)**.

### Camera calibration & kinematics analysis

The head, body, wing kinematics, and perch movements of 400 bird landing trials (100 per bird) on a moving (300 trials) and static (100 trials) perch acquired by the IDT cameras were digitized using DLTdv6 program (17) (Mathworks, USA) using manual and auto tracking features (18). Stereo 3D calibration of the cameras was carried out using a calibration wand to reconstruct the direct linear transformation (DLT) coefficients to determine the 3D coordinates (19). The two cameras provided a calibrated volume of 0.648 m3 (Fig. 1D). The average value of the re-projection error was 0.22 pixels. The tracked kinematics data followed the method described by Warrick et al (18). The following variables were computed from the high-speed cameras: (a) the bird’s head, body, and wing position; (b) bird and perch velocity; (c) angular position and phase of the perch; (d) the bird’s pitch angle relative to the perch; and (e) maximum wingspan. The bird’s pitch angle pre- and post-landing (measured when post-landing pitch torque reached an initial minimum value) was defined with respect to horizontal (Fig. 2A). The bird’s landing time was determined both when the bird’s feet made contact with the perch in preparation for landing (***t* _CONTACT_**), computed from the time when Fx and Fz resultant crossed a threshold value, set at 10% of peak force and when the bird’s feet grasped the perch (***t* LAND**), computed when the Euclidian distance between the bird relative to the perch reached a minimum value.

The bird’s landing phase (at ***t* LAND**) was computed from the perch’s angular motion and categorized into four general zones – zone 1: 0 to +¼ π and +¾ π to π (SDs; Fig. 2C); zone 2: π to +⁵⁄₄π and +⁷⁄₄ π to 2π (ODs; Fig. 2D); zone 3: +¼π to +¾π (SDf; Fig. 2E); zone 4: +⁵⁄₄π to +⁷⁄₄π (ODf; Fig. 2F) which were used to test the four hypothesized landing conditions.

### Perch torque and force recordings

Perch landing forces in the horizontal (Fx) and vertical (Fz) directions and torque about the perch (T_Y_) were acquired from the ATI force/torque sensor using a USB DAQ (USB 6255, National Instruments, USA) at 10 kHz in LabVIEW (National Instruments, USA) running a custom written LabVIEW script. (Suppl Fig. 1 A, B and C show representative recordings for different landing conditions). Both IDT cameras and the ATI load-cell were electronically synchronized using the camera’s trigger pulse.

### Statistical analysis of data

A multimodal Gaussian curve fitting algorithm (Mathworks, USA) was used to determine the frequency distribution (number of landings) of the birds’ landing phases relative to the perch’s motion (Suppl. Fig. 2). Two statistical tests were carried out to evaluate the modality of the observed frequency distribution of landing phases: Hartigan’s dip statistic (HDS) (20), and Gaussian goodness of fit (Mathworks, USA). A one-way analysis of variance (Mathworks, USA) (21), followed by a multiple comparison statistical analysis (Mathworks, USA) (22) was carried out to evaluate statistically significant differences in landing force and torque values (post landing), and body pitch angle (post landing) across the four landing conditions in comparison to stationary perch (control) landings (Fig. 2), as well as across the three perch swing frequencies versus control. Linear regression (Mathworks, USA) was carried to evaluate the correlation between the peak resultant landing force (F_R_) versus relative approach speed (XZ) between the bird and the perch (Suppl. Fig. 3).

## Results and Discussion

Representative examples of changes in bird pitch angle, wingspan movements, landing forces, and pitch torque in relation to angular position of the perch are summarized in Supplementary Figure 1 for three different landing conditions (So, SDf and ODf). For landings to a stationary perch (So) and same direction landings (SDf) no wing movements were observed post-landing, but for opposite direction landings when the perch was moving fast mid-swing (ODf), some wing movements are observed later in post-landing, presumably to help stabilize the bird on the perch. Across all landing conditions the birds approached the perch by pitching up to a consistent angle (81.9 ± 9.4o SD, ± 0.37o SEM), and subsequently pitched head-down post-landing. Angular body pitch of the bird post-landing was inversely correlated with the magnitude and direction of pitch torque exerted by the feet on the perch, such that a more negative landing pitch torque was associated with reduced head-down pitch rotation of the bird. This was the case both for landings across conditions (Fig. 5A vs 5B), as well as landings at higher perch frequencies (Fig. 5C vs 5D). Most commonly at landing (***t* _CONTACT_**), a clear surge in decelerating (negative) Fx was observed, together with modest increases in Fz (see supplemental Movies 1-4). Considerable variation in pitch torque (T_Y_) was observed post-landing, until the bird achieved a stable position on the perch.

### Landing phase timing

Consistent with our primary hypothesis, lovebirds timed their landings in a majority of trials (51.3%), across the three swing frequencies, to when the perch was approaching either extreme of its motion with its velocity nearing zero (27.5% in the same direction as the bird’s approach – SD_S_, and 23.8% in the opposite direction to the bird’s approach – ODs; Fig. 3). Less commonly, lovebirds landed when the perch was moving at higher velocity either toward the bird’s approach (12.3%, ODf) or in the same direction as the bird’s approach (SDf, 11.5%), but these landings were broadly distributed and less frequent over the middle third of the perch’s swing in both directions. The bias in landing phase for both same and opposite direction landings to ∼ ¹⁷⁄₂₀ π to ²³⁄₂₀ π (Fig. 3) likely reflects additional looming cues produced by the corridor’s end wall section, given that looming cues are known to influence flight behavior (23, 24), affecting flight paths due to optic flow cues (25–27) and guiding landing to a stationary perch (1, 3, 4, 13, 28).

**Figure 3.**
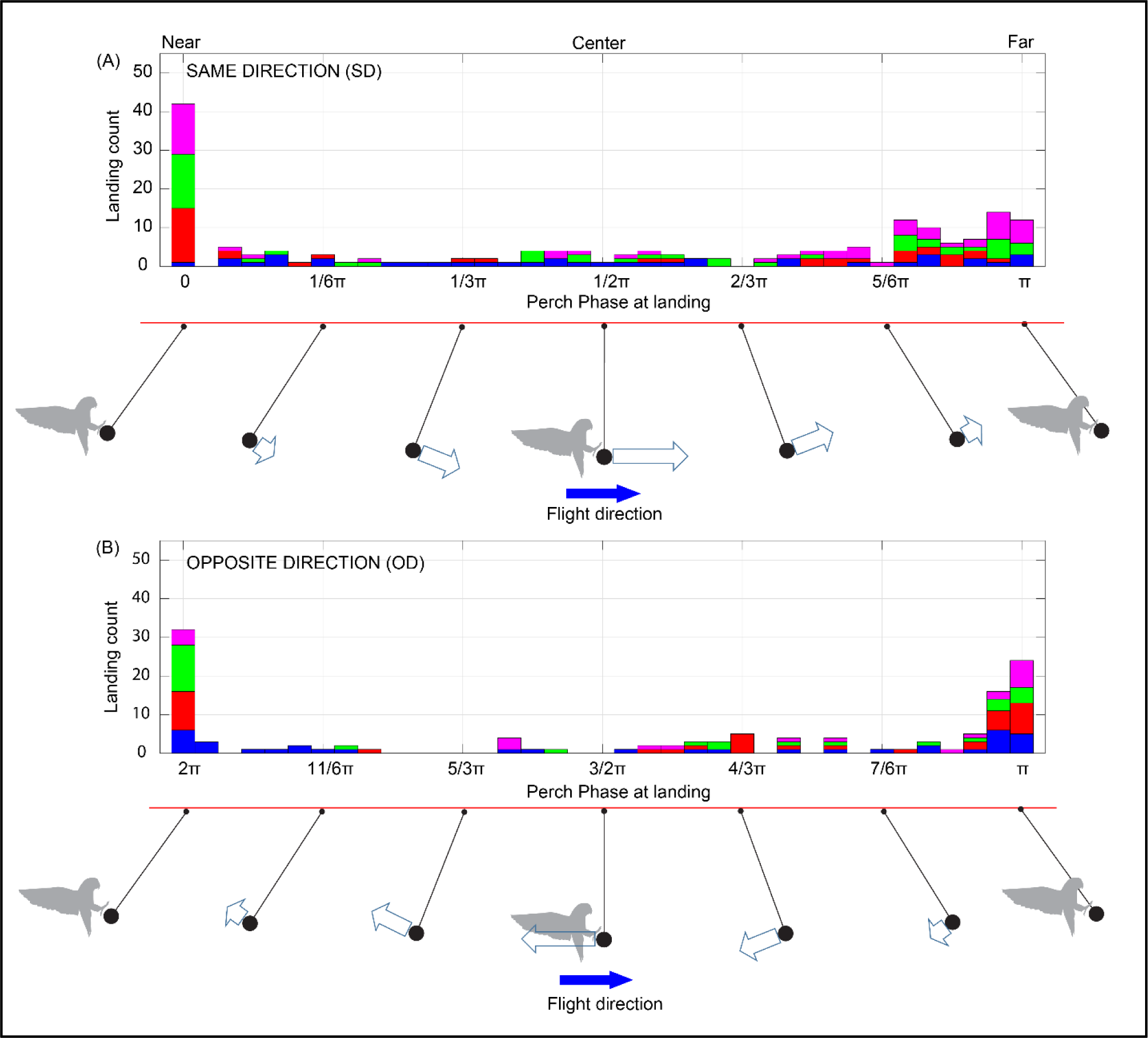
Lovebird landings on a swinging perch are strongly biased to the extremes of motion, when perch velocity is near zero. A histogram of bird landings (**A**) when the perch was moving in the same direction (**SD**) (n: 176) as the bird, and (**B**) when the perch was moving in the opposite direction to the bird (**OD**) (n: 124). Individual birds are color coded. All birds showed a preference to land when the perch was moving slowly at the extreme ends of its oscillation (0, π or 2π), approaching zero velocity. Due to the broader distribution of landing between ∼ ¹¹⁄₂₀ π to π across all birds for **SD** (50%) and π to + ¹³⁄₁₀ π **OD** (62.9%) landings, a higher percentage of landings occurred during this phase range of the perch’s motion. As a consequence, there was a higher percentage of landings when the perch had swung towards the far end of the corridor, suggesting that looming cues induced by the closed end of the corridor (23, 24) may have biased landings to this phase of the perch’s motion. (See Suppl. Fig. 2).

We carried out two statistical tests to assess the modality of the lovebirds’ landing phase with respect to the swinging perch (Table 1). Hartigan’s dip statistic (HDS) analysis indicates support for same direction (Dip = 0.1159, *p* < 0.0001) and opposite direction (Dip = 0.1290, *p* < 0.0001) bimodal distributions. A second analysis of the frequency distribution of landing phase counts using a ‘Gaussian 3’ curve fit (Mathworks, USA) gives a ‘goodness of fit’ R2 values of 0.9199, SSE (Sum of Squared Errors): 149.1 (*p* < 0.0001) (same direction) and 0.8933, SSE: 157.7 (*p* < 0.0001) (opposite direction), indicating a strong bimodal and weak multimodal distribution (Suppl. Fig. 2).

### Landing forces and pitch torque in relation to landing kinematics

Our hypothesis that birds would time their landings to minimize landing force and pitch torque was partly supported, as we found that landing force (Fx & F_R_) was greater in opposite direction versus same direction landings (Fig. 4), consistent with greater resultant landing force associated with the bird’s relative approach speed (Suppl. Fig. 3). However, no clear pattern of reduced landing pitch torque emerged (Fig. 5).

**Figure 4.**
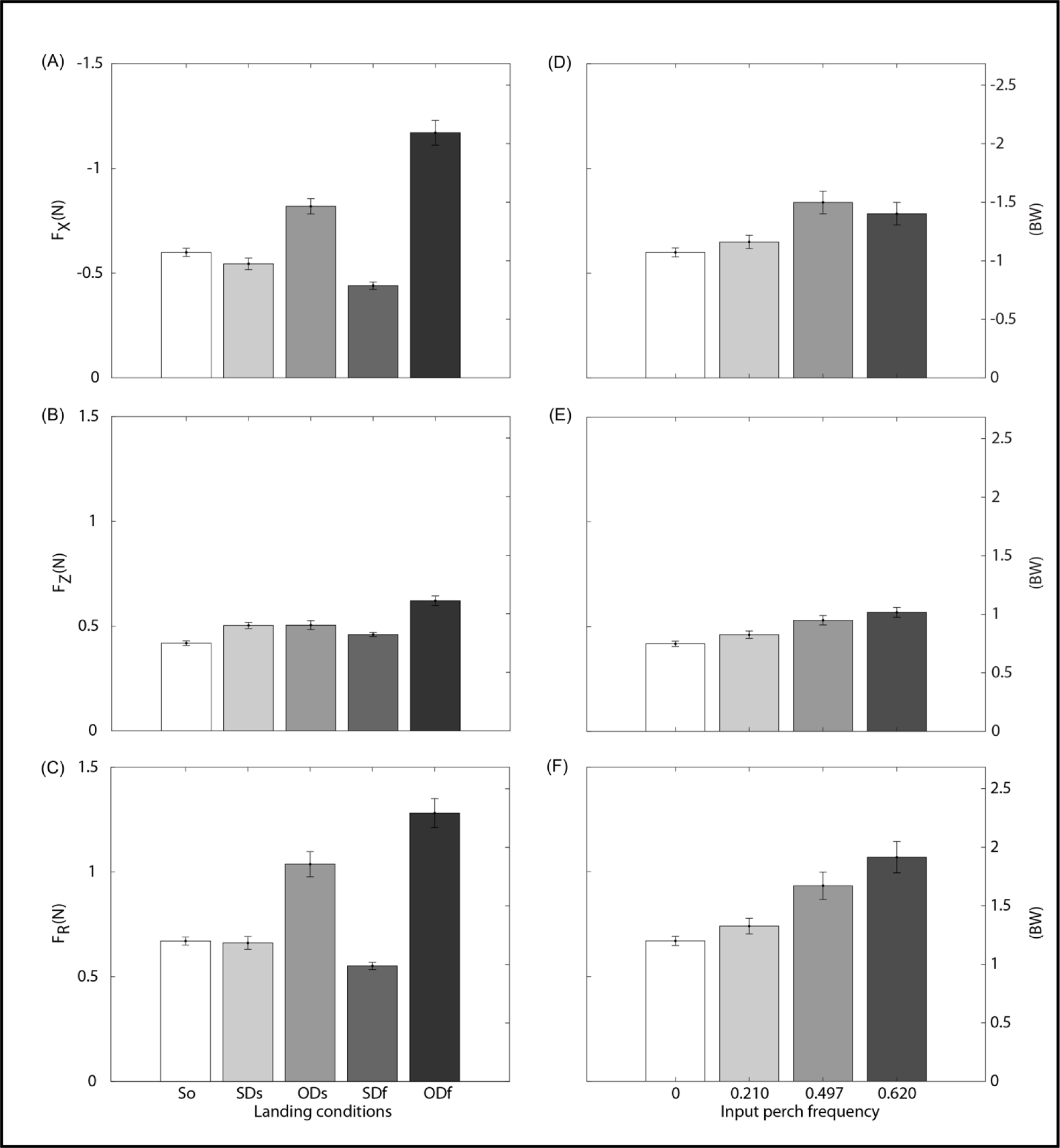
Greater landing force occurred for opposite direction perch landings and at higher perch frequency. Histograms showing the mean (A) horizontal component (**Fx**), (B) vertical component (**Fz**) and (C) resultant (**F_R_**) landing force for various landing conditions: stationary control (**So**), same direction slow (**SDs**), opposite direction slow (**ODs**), same direction fast (**SDf**), and opposite direction fast (**ODf**). (D), (E) and (F) show landing force patterns in relation to perch frequency. Error bars denote ± SEM. One way ANOVA and multiple comparison analysis (see Methods) show that landing forces significantly differ between most landing and frequency conditions, being greatest for **ODf** and the highest frequency conditions. Vertical landing forces averaged ∼50% of horizontal forces (Data. N=4; n = 400; see Tables 2 and 3).

**Figure 5.**
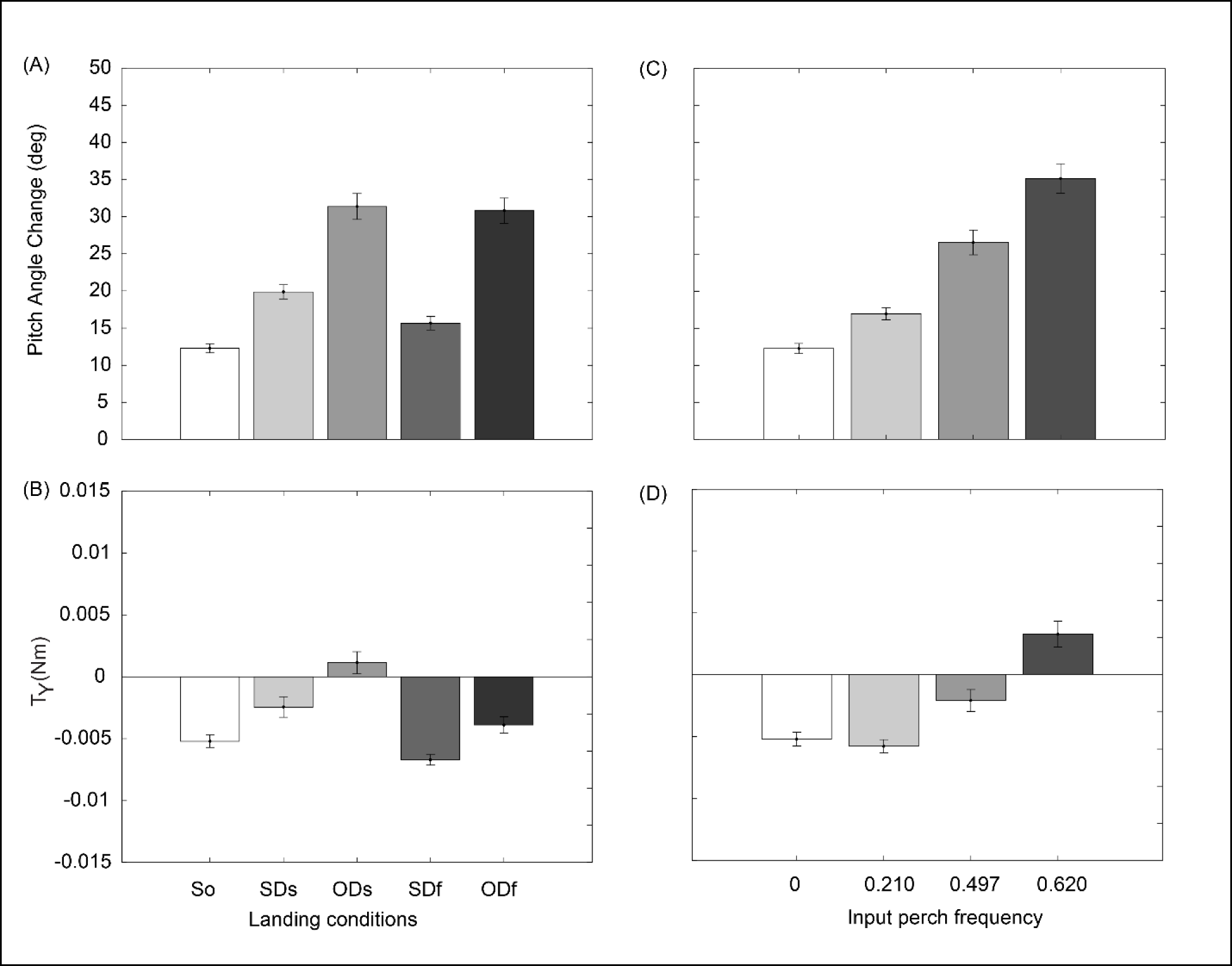
Post-landing pitch angle change is correlated with maximum pitch torque *(T_Y_)* for different landing conditions and perch frequencies. Histograms of **(A)** change in body pitch angle relative to the perch (+ head down) measured from the onset of landing to the time of **(B)** maximum post-landing torque (**T_Y_**). Pitch angle change varies directly with landing torque (R = 0.57696, *p* < 0.0001, R2=0.333; least-squares regression), with both exhibiting variable patterns across landing conditions. **(C)** Body pitch angle change and **(D)** maximum pitch torque increased with increasing perch frequency and again were correlated (R = 0.57676, *p* < 0.0001, R2=0.333; least squares regression). Error bars represent ± SEM. One way ANOVA and multiple comparison analysis (see Methods) show that landing torque and body pitch angle change significantly differ between most landing and frequency conditions. (Data. N = 4; n = 400; see Tables 4 and 5).

**Table 2:**
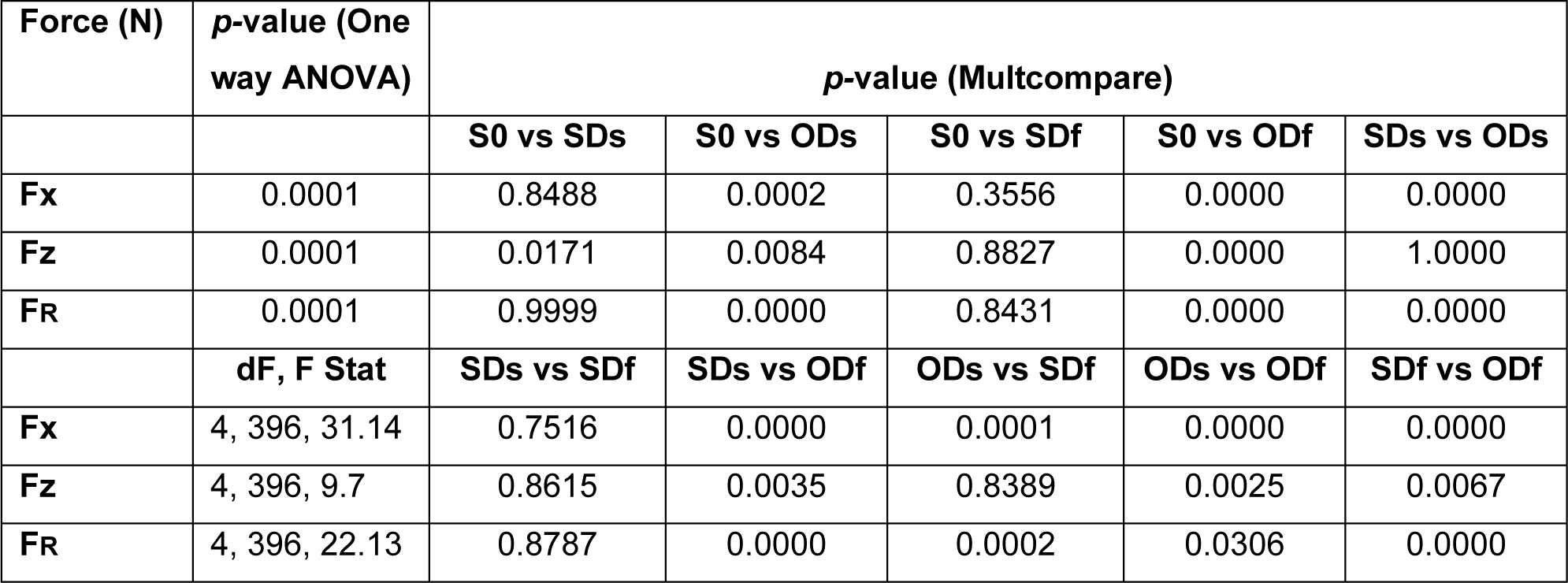
Results of *p*-values, dF and F-statistics of ANOVA (One way) and Multcompare for Fx, Fz and FR across different landing conditions. (n= 400).

| Force (N) | p-value (One way ANOVA) | p-value (Multcompare) |  |  |  |  |
| --- | --- | --- | --- | --- | --- | --- |
|  |  | S0 vs SDs | S0 vs ODs | S0 vs SDf | S0 vs ODf | SDs vs ODs |
| **F<sub>x</sub>** | 0.0001 | 0.8488 | 0.0002 | 0.3556 | 0.0000 | 0.0000 |
| **F<sub>z</sub>** | 0.0001 | 0.0171 | 0.0084 | 0.8827 | 0.0000 | 1.0000 |
| **F<sub>R</sub>** | 0.0001 | 0.9999 | 0.0000 | 0.8431 | 0.0000 | 0.0000 |
|  | dF, F Stat | SDs vs SDf | SDs vs ODf | ODs vs SDf | ODs vs ODf | SDf vs ODf |
| **F<sub>x</sub>** | 4, 396, 31.14 | 0.7516 | 0.0000 | 0.0001 | 0.0000 | 0.0000 |
| **F<sub>z</sub>** | 4, 396, 9.7 | 0.8615 | 0.0035 | 0.8389 | 0.0025 | 0.0067 |
| **F<sub>R</sub>** | 4, 396, 22.13 | 0.8787 | 0.0000 | 0.0002 | 0.0306 | 0.0000 |

**Table 3:** Results of *p-*values, dF and F-statistics of ANOVA (One way) and Multcompare for Fx, Fz and FR across different input perch frequencies. (n= 400).

| Force (N) | *p*-value (One way ANOVA) | *p*-value (Multcompare) |  |  |  |  |  |
| --- | --- | --- | --- | --- | --- | --- | --- |
|  |  | Cont vs<br>0.2 Hz | Cont vs<br>0.46 Hz | Cont vs<br>0.6 Hz | 0.2 vs<br>0.46 Hz | 0.2 vs<br>0.6 Hz | 0.46 vs<br>0.6 Hz |
| **F<sub>x</sub>** | 0.0001 | 0.8430 | 0.0004 | 0.0117 | 0.0095 | 0.1124 | 0.8112 |
| **F<sub>z</sub>** | 0.0001 | 0.3958 | 0.0003 | 0.0000 | 0.0584 | 0.0006 | 0.5211 |
| **F<sub>R</sub>** | 0.0001 | 0.7930 | 0.0032 | 0.0000 | 0.0561 | 0.0001 | 0.2839 |
|  | **dF, F Stat** |  |  |  |  |  |  |
| **F<sub>x</sub>** | 3, 396, 6.88 |  |  |  |  |  |  |
| **F<sub>z</sub>** | 3, 396, 11.99 |  |  |  |  |  |  |
| **F<sub>R</sub>** | 3,396, 11.35 |  |  |  |  |  |  |

**Table 4:**
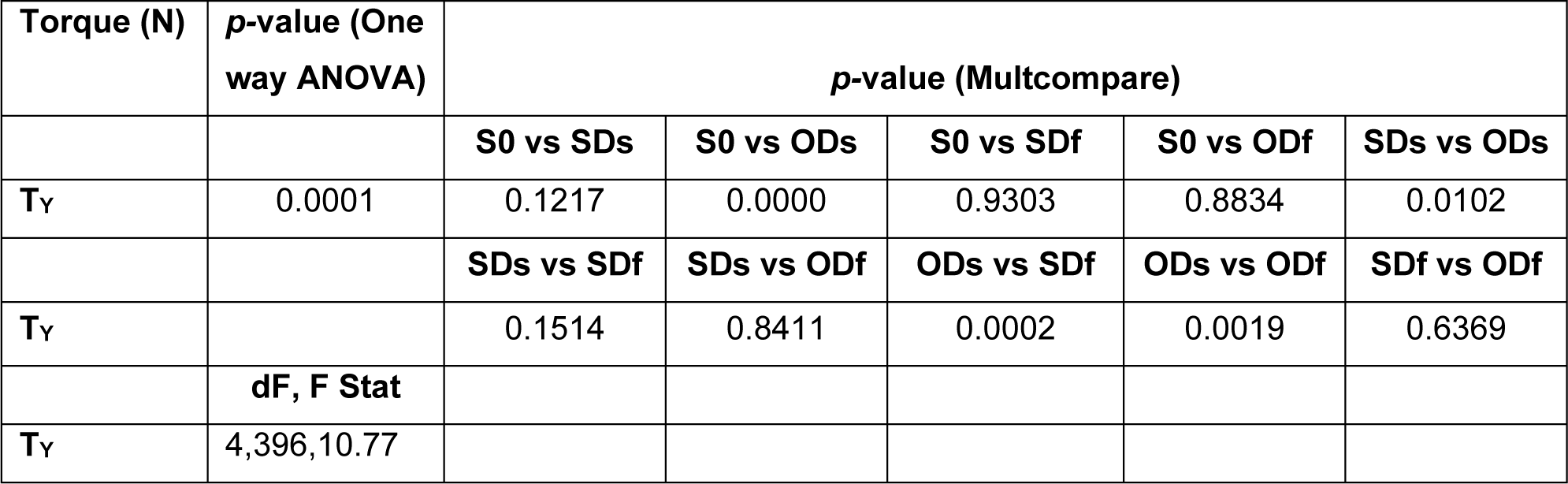
Results of *p*-values of ANOVA (One way) and Multcompare for post-landing pitch torque (TY) across different landing conditions. (n= 400).

**Table 5:** Results of *p*-values of ANOVA (One way) and Multcompare for post-landing pitch torque (TY) across different input perch frequencies. (n= 400).

| Torque<br>(N m) | *p*-value (One<br>way ANOVA) | *p*-value (Multcompare) |  |  |  |  |  |
| --- | --- | --- | --- | --- | --- | --- | --- |
|  |  | Control<br>vs 0.2 Hz | Control<br>vs 0.46 Hz | Control<br>vs 0.6 Hz | 0.2 Hz vs<br>0.46 Hz | 0.2 Hz vs<br>0.6 Hz | 0.46 Hz<br>vs 0.6 Hz |
| **T<sub>Y</sub>** | 0.0001 | 0.9557 | 0.0259 | 0.0000 | 0.0050 | 0.0000 | 0.0000 |
|  | **dF, F Stat** |  |  |  |  |  |  |
| **T<sub>Y</sub>** | 3,396, 27.76 |  |  |  |  |  |  |

**Table 6:** Results of *p*-values of ANOVA (One way) and Multcompare for post-landing body pitch angle change across different landing conditions. (n= 400).

|  | *p*-value<br>(One way<br>ANOVA) | *p*-value (Multcompare) |  |  |  |  |
| --- | --- | --- | --- | --- | --- | --- |
|  |  | S0 vs SDs | S0 vs ODs | S0 vs SDf | S0 vs ODf | SDs vs ODs |
| **Pitch Angle**<br>**Change** | 0.0001 | 0.0020 | 0.0000 | 0.8423 | 0.0000 | 0.0000 |
|  |  | SDs vs SDf | SDs vs ODf | ODs vs SDf | ODs vs ODf | SDf vs ODf |
| **Pitch Angle**<br>**Change** |  | 0.7093 | 0.0001 | 0.0000 | 0.9994 | 0.0002 |
|  | **dF, F Stat** |  |  |  |  |  |
| **Pitch Angle**<br>**Change** | 4,396, 30.34 |  |  |  |  |  |

**Table 7:** Results of *p*-values of ANOVA (One way) and Multcompare for post-landing body pitch angle change across different perch frequencies. (n= 400).

|  | **p-value**<br>**(One way**<br>**ANOVA)** | **p-value (Multcompare)** |  |  |  |  |  |
| --- | --- | --- | --- | --- | --- | --- | --- |
|  |  | **Control**<br>**vs 0.2 Hz** | **Control**<br>**vs 0.46 Hz** | **Control**<br>**vs 0.6 Hz** | **0.2 Hz vs**<br>**0.46 Hz** | **0.2 Hz vs**<br>**0.6 Hz** | **0.46 Hz vs**<br>**0.6 Hz** |
| **Pitch Angle**<br>**Change** | 0.0001 | 0.0853 | 0.0000 | 0 | 0.0000 | 0.0000 | 0.0001 |
|  | **dF, F Stat** |  |  |  |  |  |  |
| **Pitch Angle**<br>**Change** | 3,396, 52.84 |  |  |  |  |  |  |

Horizontal landing forces exceeded vertical landing forces across all landing conditions (Fig. 4). Horizontal forces (Fx Fig. 4A) were lowest when the perch was stationary (−0.59 N) or moving in the same direction at slow speed and fast speeds −0.54 & −0.44 N, respectively (= −1.02 & −0.83 BW), in support of our hypothesis for conditions So and SDs (Fig. 2A & B), but not for SDf (Fig. 2E). Fx was greatest for ODs and ODf landing conditions (−0.82 & −1.17 N, respectively; = −1.44 & −2.21 BW); increasing by two-fold (*p*<0.05, multicompare analysis, Table: 2) relative to the So and SDs conditions, in support of our hypothesized ODf landing condition (Fig. 2E). Greater horizontal landing (Fx) forces were recorded at higher perch swing frequencies (Fig. 4B), being - 0.84 N at 0.467 Hz. (*F*_3, 396_: 6.88, *p* < 0.0001; Table 3).

Counter to our hypotheses for landing pitch torque (T_Y_) magnitude and direction, the lovebirds exerted a negative (head-up) pitch torque across nearly all landing conditions and perch frequencies (Fig. 5B & D; (exceptions being the low magnitude positive pitch torque during ODs and highest frequency landings)). Positive pitch torque was not observed for opposite direction landings (Fig. 2D & F) nor was pitch torque greater for landing when the perch was moving fast (middle of swing) versus slow (ends of swing; Fig. 2 C, D vs E, F). As expected, the magnitude of pitch torque varied strongly with respect to the angular motion of the bird on the perch post-landing (Fig. 5, Suppl. Fig 3, and Suppl. Movies 1-4) across landing conditions (R2= 0.333, *p* <0.0001) and differing perch frequencies (R2= 0.333, *p* <0.0001). Lovebirds also exhibited greater head-down pitch rotation during opposite direction landings compared with stationary and same direction landings (Fig. 5A; *F*_4,396_: 10.77, *p* <0.0001), as summarized in table 4.

The 1.46-fold greater horizontal versus vertical landing force observed across all trials reflects the shallow landing approach (COM: −13.2 ± 3.0o relative to horizontal (Suppl. Fig. 5D) that the lovebirds adopted to land (Mobius camera close-up video recordings; Suppl. Movies 1-4). By approaching with a shallow (negative) angle the lovebirds reduce the orthogonal distance of their COM to the perch (3.61 ± 0.21 cm) at landing and thus, the magnitude of pitch torque subsequently produced about the perch. However, this requires them to elevate their feet by upward pitch of their trunk with their ankle and knee initially extended prior to landing, to achieve a stable foot landing angle of 56.9 ± 2.8o (Suppl. Fig. 5). By pitching their body up to 82 degrees at landing, the lovebirds’ achieve a nearly horizontal stroke plane with their final wingbeats, with the wing chord aligned close to vertical when the wings were elevated. Although the low temporal resolution (60 Hz) and restricted close-up field of view of the Mobius videos (see Suppl. Movies 1-4) prevented an accurate measure of the wings’ stroke plane kinematics, this orientation of the wings likely provided a decelerating aerodynamic force (not measured) to slow the bird’s approach, in combination with the horizontal landing force exerted at the perch to arrest their momentum. Following perch contact, the lovebirds then absorbed the decelerating landing force exerted on the perch by allowing their hind limb joints (primarily their ankles and knees) to flex relative to their trunk (Suppl. Fig. 5).

As noted above, the bird’s body pitch angle during aerial approach to initiate foot contact with the perch was remarkably consistent for the different landing conditions with respect to phase (81.9 ± 0.37o SEM) and perch frequency (81.9 ± 0.46o SEM). However, the change in body pitch angle post-landing (Fig. 5A & C) generally varied inversely with respect to the magnitude and direction of peak landing pitch torque (Fig. 5B & D). This apparent discordance between observed body pitch rotation post landing and landing torque ignores the fact that birds do not land rigidly, as represented in Fig. 2, but can modulate angular changes (flexion) of their hind limb joints relative to their trunk, as well as the trajectory of their center of mass relative to the perch at landing (Suppl. Fig. 5). Such adjustments likely enable them to reduce the magnitude of pitch torque, effectively absorb their body’s kinetic energy when decelerating, and control their body pitch post landing to achieve stable landings.

Our observations reported here for peach-faced lovebirds are the first to examine biomechanical and kinematic details of avian landing on a swinging perch. Prior work (5) examined the visual cues that lovebirds use to turn and locate a hanging perch that was stationary but allowed to move in response to the bird’s take-off. Their study did not examine how lovebirds timed their return landing to the motion of the perch, or how the birds stabilized upon landing. Other studies have examined the takeoff versus landing forces (29, 30) from fixed perches, as well as the kinematics and aerodynamics (31) of birds landing on stationary perches. Consistent with our main hypothesis, we found that lovebirds favored timing their landings to when the perch’s velocity was minimal (at the beginning and end of its swing). As has been recently shown (14), closely related Pacific parrotlets exhibit excellent gripping performance when contacting stationary perches of varying texture and diameter. Consequently, it may not be surprising that the lovebirds exhibited substantial variation in landing torque after gripping the perch to help them stabilize their body as the perch continued to move.

The lovebirds’ uniform increase in body pitch to a consistent angle of 82o relative to horizontal as they approached to land on a moving as well as a stationary perch is consistent with observations for other flying and gliding animals (7, 10, 11). However, the body pitch angle of 82o adopted by lovebirds across all landing conditions exceeded the body angle adopted by diamond doves (61o) and zebra finches (59o) when landing on a stationary platform (9). This likely reflects the more inclined vertical approach that zebra finches and diamond doves adopted when landing, compared with the shallower horizontal landing approach of the lovebirds. By pitching up the birds shift their wings’ stroke plane to a more horizontal orientation, which has been shown to rotate the angle of induced downwash produced by the wings of diamond doves and zebra finches (9), and to shift the components of lift and drag forces that Pacific parrotlets generate (32) when landing on a stationary perch. As a result, both aerodynamic lift and drag contribute to slowing the bird’s approach velocity, allowing the bird to land with reduced perch reaction forces and resulting pitch torque, as observed here for lovebirds landing on a swinging perch.

Visual cues used by birds and insects to locate and target landing sites (1–4) or to dock at flowers while hovering to feed on nectar (33, 34) have mainly focused on stationary targets; but see Sponberg et al. (2015) (34) for moths tracking moving flowers under low light conditions. The visual cues that appear most relevant to birds (as well as flying insects, (35, 36) to target landing sites and avoid obstacles are the estimated time-to-collision (‘tau’-based cue) (2, 37) or the relative size of retinal image motion (RREV) based on optic flow cues (1, 36). In their study of lovebirds landing to a suspended perch that was allowed to swing after the bird’s take-off, Kress et al, (5) found that the inferred retinal size of the perch appeared to provide the clearest visual cue to target leg extension and landing at short range rather than tau or RREV; which may operate to gauge distance and time landing or obstacle avoidance at longer distances. Given these varying results reported to date based on how flying animals target stationary versus moving perches as landing sites, additional analysis of visual cues used by birds to time their landing relative to the phase of perch motion and perch location to achieve robustly stable and safe landing, is warranted.

### Future directions and applications

Our observations, which are the first to investigate the biomechanics and kinematics of birds landing on a moving target, show that this behavior is a precisely controlled and timed event that favors landing when the landing target is moving most slowly. More importantly, by adopting a horizontal approach trajectory to the perch, the lovebirds are not only able to decelerate their landing momentum but also reduce the likelihood of landing errors when contacting and gripping to stabilize on the perch. This also most likely simplifies the visual cues needed for stable and safe landing.

In this context our results have bioinspired relevance for improving the flight and landing control of unmanned aerial vehicles (UAV) on a moving platform. Biologically inspired strategies have been used to control the autonomous flight and landing of a fixed wing drone (38), and have explored passive mechanisms by which a helicopter or a quadcopter drone can be made to land and grab on a branch of a tree (39), remaining stably in place by means of its passive weight, even when experiencing a wind gust (40). However, no previous studies have been carried out in which biological inspired principles were used to guide a drone or fixed wing UAV to land on a moving branch of a tree. Results from the present study may help to guide improved approaches by applying bioinspired principles based on how flying animals target and land robustly on moving perches.

**Supplementary Movie 1.**

Landing condition: SDs – shows modest negative (head-up) pitch torque at landing with similar Fx and Fz landing forces, followed by positive (head-down) pitch torque and greater Fx reaction force.

**Supplementary Movie 2.**

Landing condition: SDf – shows initial relatively large negative (head-up) pitch torque at landing with greater Fx versus Fz landing force, followed by modest negative pitch torque and landing forces as the bird stabilizes on the perch.

**Supplementary Movie 3.**

Landing condition: SDs – shows initially small negative (head-up) pitch torque at landing, followed by positive (head-down) pitch torque as the bird stabilizes on the perch. Fx forces generally exceed Fz forces throughout the landing.

**Supplementary Movie 4.**

Landing condition: ODs – shows initial negative (head-up) pitch torque at landing, followed by moderate positive pitch torque, with Fx forces that exceed Fz forces throughout the landing.

## Acknowledgements

We would like to thank Ms. Lilly Lu (data collection); Dr. Ravi Mane Tanaji & Dr. Marcus Hietanen (Matlab support); Dr. Andrew Mountcastle (3D printing). Prof. MV Srinivasan, Dr. Ivo Ros & Prof. Nicolai Konow provided critical comments on the manuscript, guidance on data analysis & figure preparation. We thank Mr. Pedro Ramirez for animal care and husbandry.

## Author contributions

**(Following CRediT taxonomy):** conceptualization: P.S.B. and A.A.B.; methodology: P.S.B. and A.A.B.; software: P.S.B.; validation: P.S.B. and A.A.B.; formal analysis: P.S.B. and A.A.B.; investigation: P.S.B. and A.A.B.; resources: P.S.B. and A.A.B.; data curation: P.S.B. and A.A.B.; writing original draft: P.S.B. and A.A.B.; writing review & editing: P.S.B. and A.A.B.; visualization: P.S.B. and A.A.B.; supervision: A.A.B.; project administration: A.A.B.; funding acquisition: A.A.B.

## Author Approvals

The authors have viewed and approved the manuscript and it has not been accepted or published elsewhere.

---

**Supplementary Figure 1.**
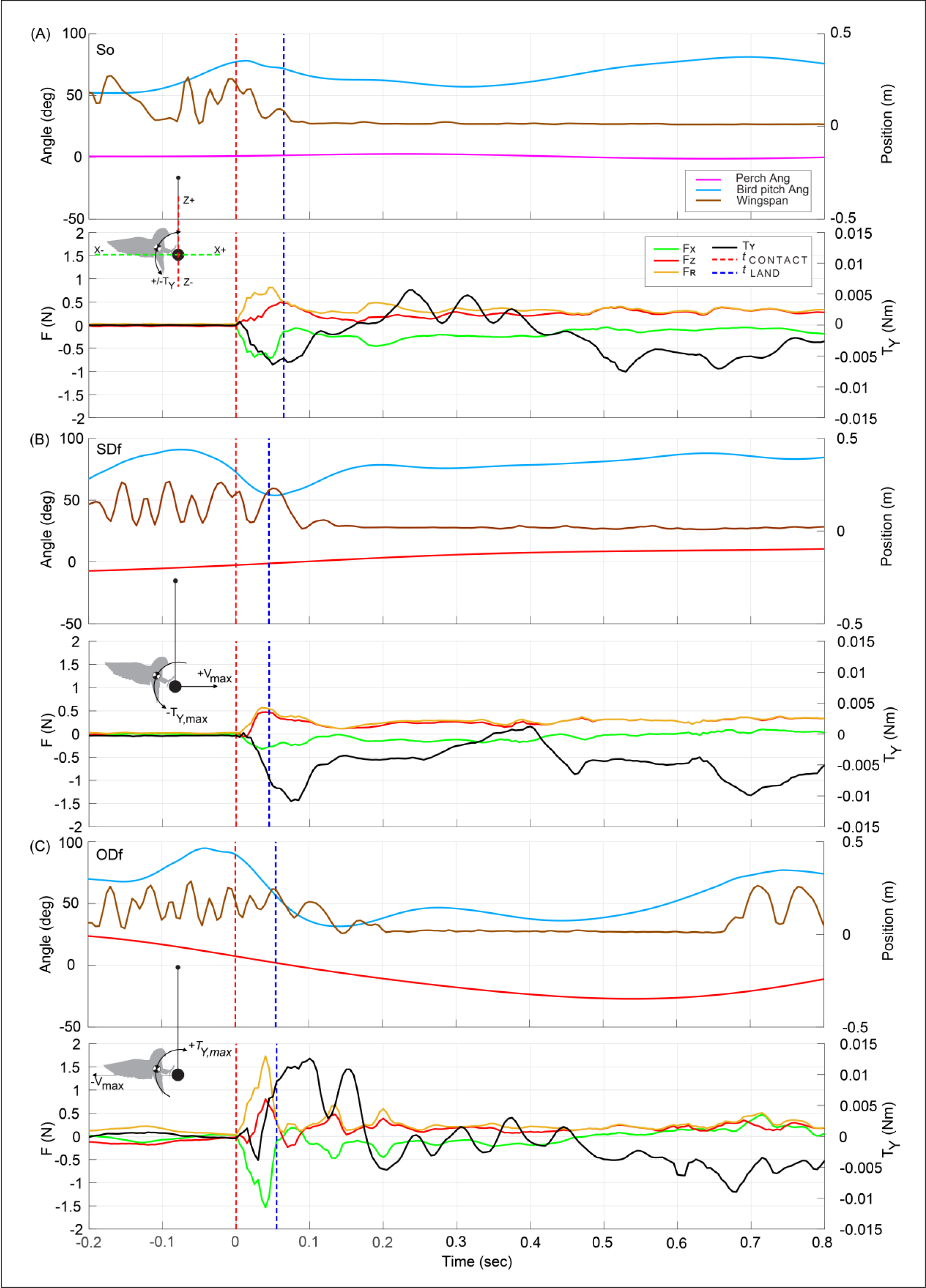
Representative examples of time-varying changes in bird pitch angle, wingspan movements, landing forces, and landing pitch torque (T_Y_) in relation to angular position of the perch for three landing conditions (So, SDf and ODf). In these example recordings, (A) during landings to a stationary perch (**So**) little or no wing movements were observed post-landing. Lovebirds pitched up to a consistent angle (on average 82o across landing conditions) to land and exhibited modest pitch motions to stabilize post-landing, associated with a rise in force (**Fx, Fz & F_R_**) and pitch torque (**T_Y_**) after ***t* CONTACT**. (B) For same direction landing (**SDf**) little to no wing movements were observed post-landing, with the bird pitching up (91°) to land in this example, with a more substantial rise pitch torque after foot contact with the perch. (C) For opposite direction landing (**ODf**) periodic wing movements were commonly observed post-landing, helping the bird to stabilize on the perch following the sharp rise in force and torque at the onset of landing. Fluctuations in pitch torque were more substantial **ODf** post-landing than for the other landing conditions. Vertical red dashed line denotes ***t* CONTACT**, and vertical blue dashed line denotes ***t* LAND.**

**Supplementary Figure 2:**
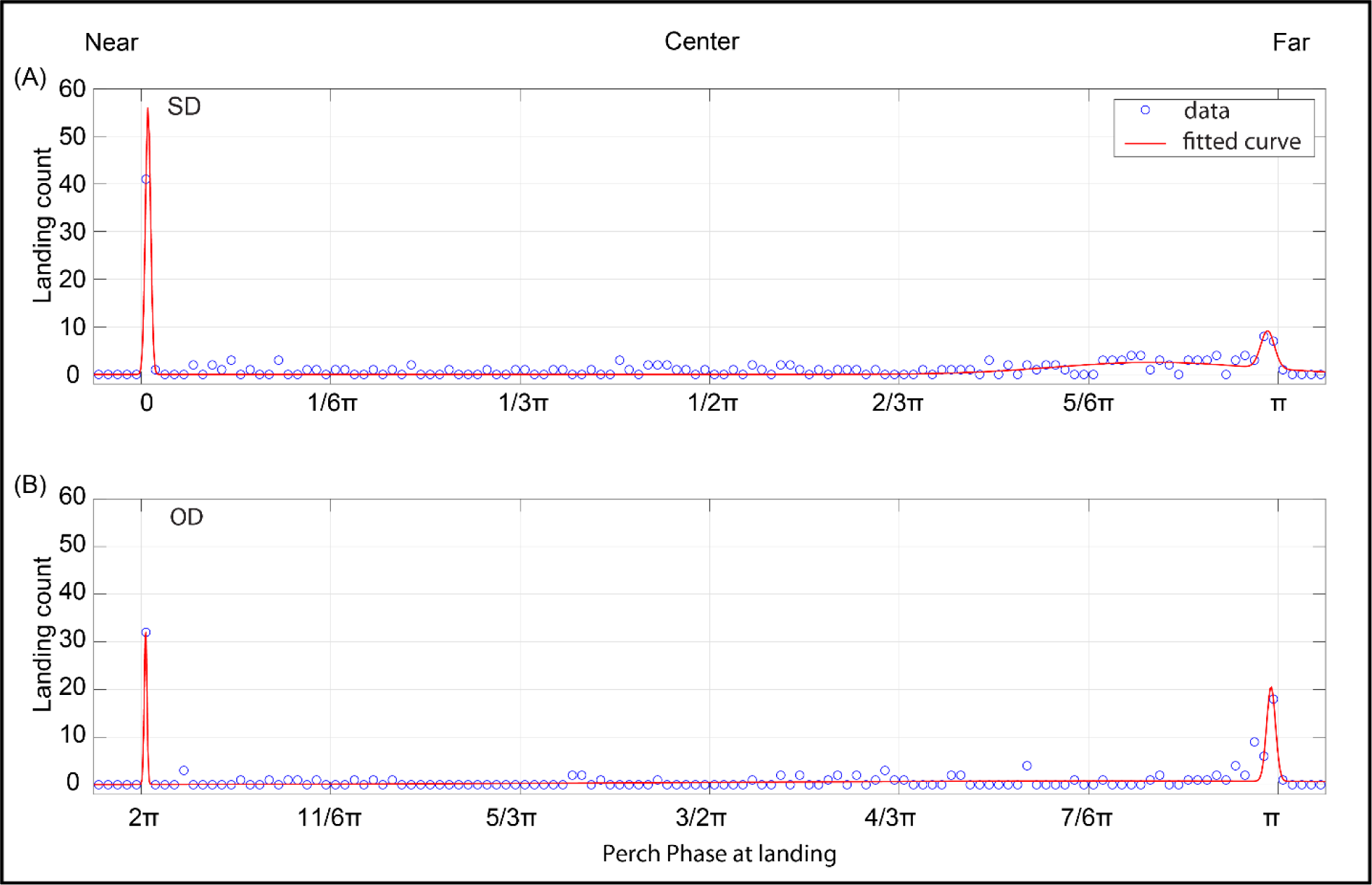
Modality patterns of lovebird landing phases on a moving perch exhibit a strongly bimodal and weakly multimodal distribution. Gaussian fit (A) **SD**: R2 = 0.9199 and (B) **OD**: R2 = 0.8933. (See Table 1). The bimodal ratio of the left (0) versus right (+π) landing peaks for **SD** landings is 0.160, and for left (2π) versus right (+π) **OD** landing peaks is 0.640. The 3rd minor landing mode centered at ∼ ¹¹⁄₁₂ π for **SD** landings is 0.0457 and none for **OD** landings.

**Supplementary Figure 3.**
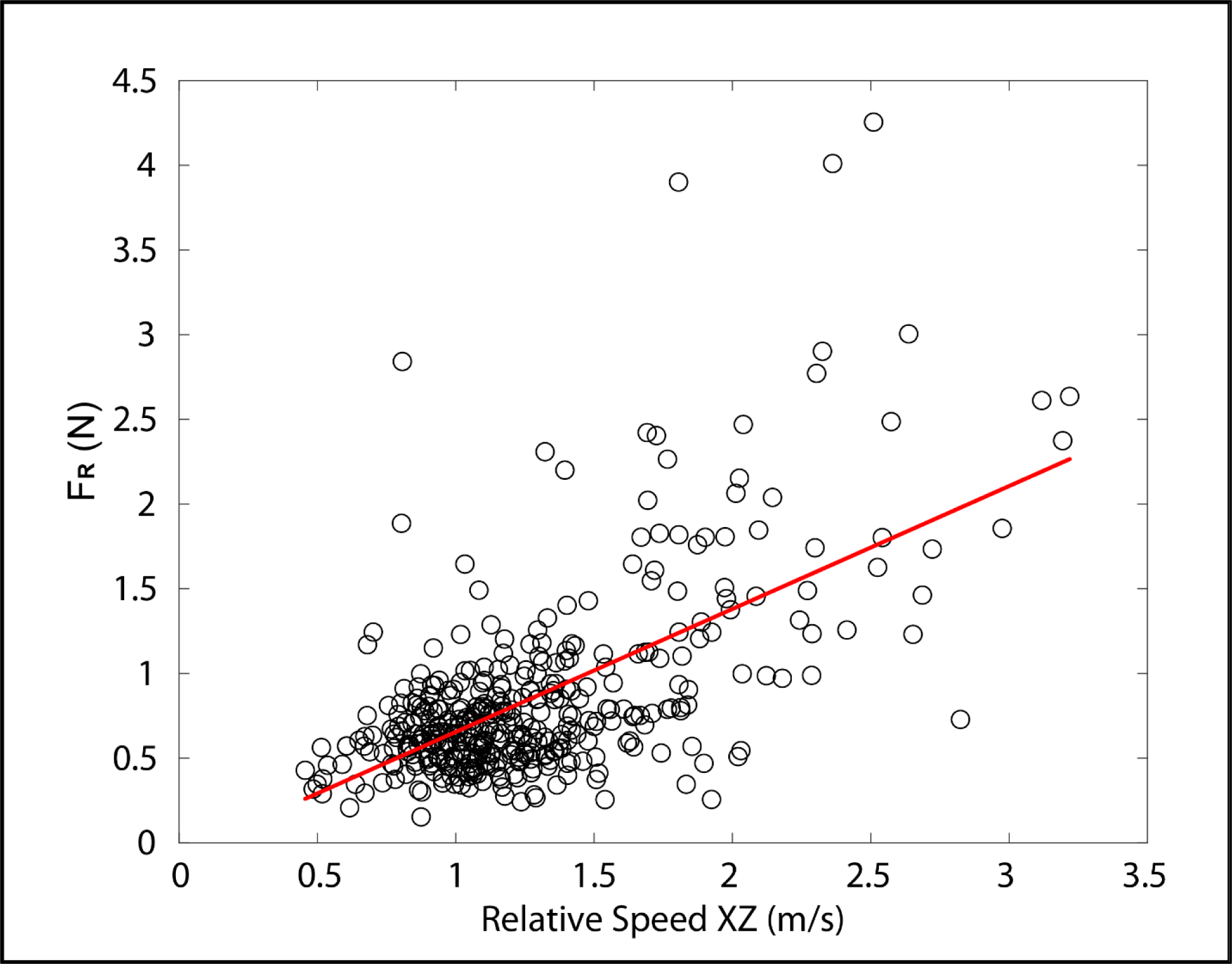
Scatter plot of peak resultant landing force versus relative approach speed across all landing conditions. A significant correlation was observed between resultant landing force (F**_R_**) versus relative approach speed (R^2^ = 0.3745, *p <* 0.0001; least-squares regression: *F*_400, 398_: 238 Root Mean Squared Error: 0.444).

**Supplementary Figure 4.**
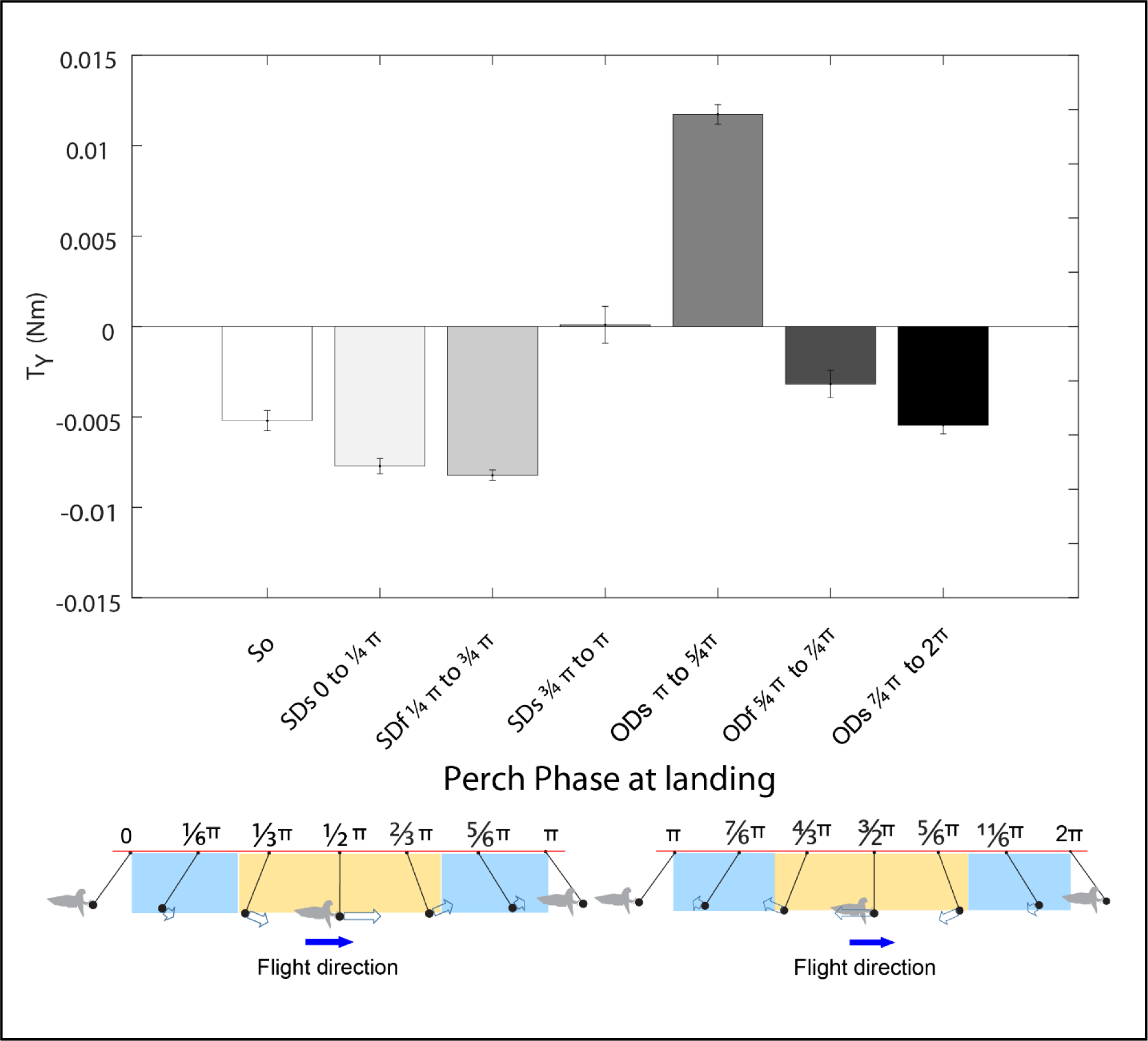
Mean ± SEM pitch torque across different perch landing phases. Stationary perch landings (**So**) resulted in a modest head-up pitch torque at landing. Same direction slow landings (**SDs**) from 0 to ¼ π (blue phase range at left, left panel) and same direction fast landings (**SDf**, ¼ π to ¾ π, yellow phase range at center, left panel) also resulted in head-up pitch torque at landing. Same direction slow landings (**SDs**) from ¾ π to π (blue phase range at right, left panel) resulted in nearly zero pitch torque. Opposite direction slow landings (**ODs**) from π to ⁵⁄₄ π (blue phase range at right, right panel) resulted in a substantial head-down pitch torque. Opposite direction fast landings (**ODf**, ⁵⁄₄ π to ⁷⁄₄ π, yellow phase range at center, right panel) and opposite direction slow landings (**ODs**) from ⁷⁄₄ π to 2 π (blue phase range to left, right panel) resulted in moderate head-up pitch torque.

**Supplementary Figure 5.**
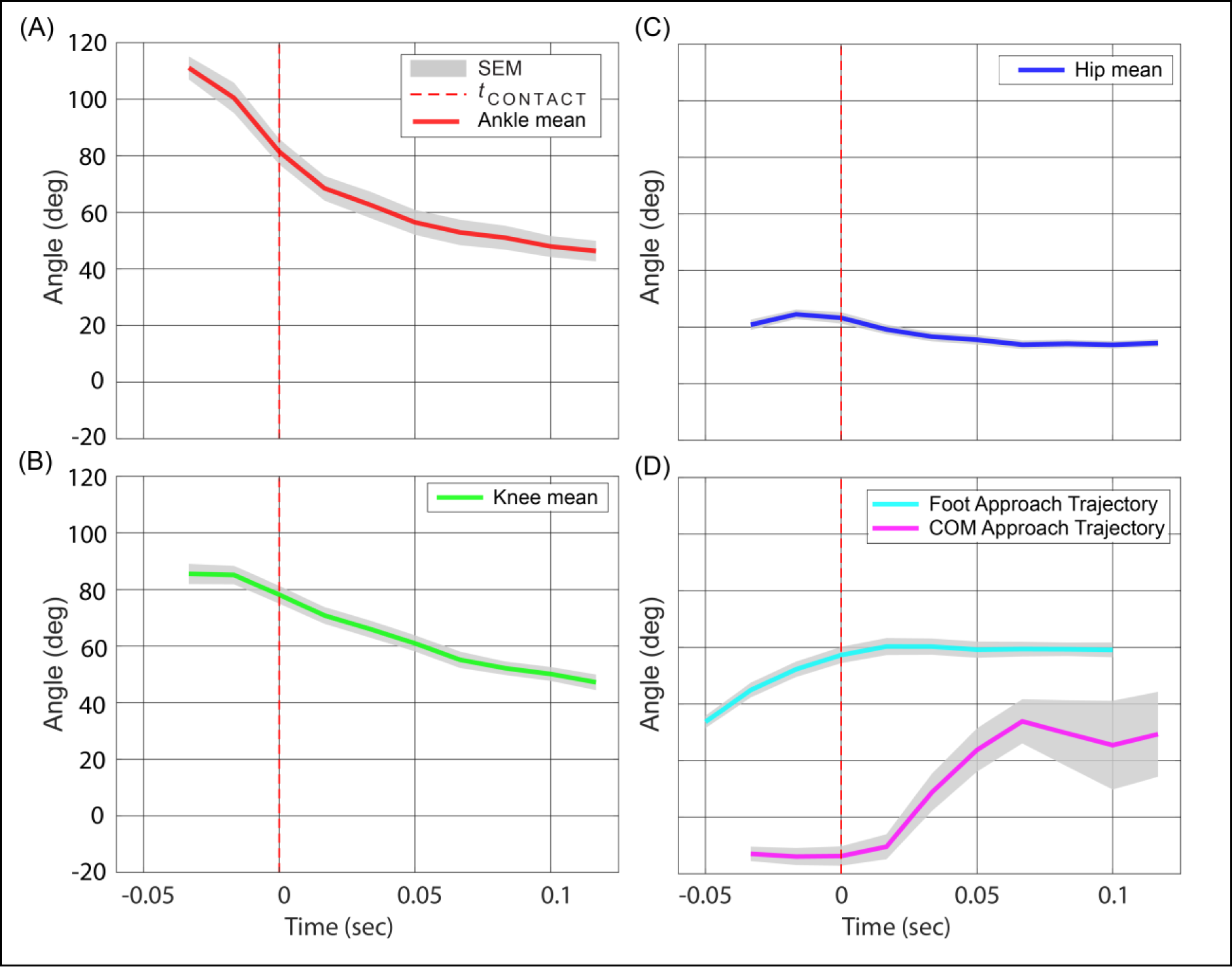
Hind limb joint joints flex and horizontal COM landing approach requires elevated foot trajectory to achieve stable landing. Changes in (A) ankle, (B) knee, (C) hip, and (D) COM (relative to horizontal) and foot approach (mid-tarsus relative to perch support) angles determined from the Mobius camera recordings at the perch. Plots show mean ± SEM angle changes over time, relative to the time of contact with the perch (vertical red dashed lines). (A & B) The ankle and knee both flex during landing (−29.2 ± 9.2o and −13.6 ± 7.4o, respectively), anticipating and then presumably absorbing the energy of impact with the perch; whereas (C) the hip remains highly flexed (mean: 21.1 ± 3.4o) with respect to the bird’s trunk throughout landing. (D) The lovebirds landed with a slightly negative COM trajectory (−13.2 ± 3.0o) as they approached the perch to land, reducing the orthogonal distance of their COM relative to the perch. This required the birds to swing their feet up by upward trunk pitch, with their ankle and knee initially extended prior to landing, to achieve a stable foot landing angle of 56.9 ± 2.8o. Data are for three of the four birds, n=36 trials (2 trials/condition).

**Supplementary table 1:** Showing <u>the</u> observed perch frequency, perch amplitude, perch angular velocity, and maximum perch speed. (n=400).

| **Serial**<br>**No** | **Perch**<br>**frequency (Hz)** | **Perch**<br>**amplitude (Deg)** | **Perch angular**<br>**velocity (Deg/sec)** | **Max perch speed**<br>**(m/sec)** |
| --- | --- | --- | --- | --- |
| 1 | 0 | 0 | 0 | 0 |
| 2 | 0.210 | 12.3 | 22.6 | 0.356 |
| 3 | 0.497 | 16.5 | 65.7 | 1.04 |
| 4 | 0.620 | 23.9 | 115 | 1.82 |

